# Electrophysiological Signature Reveals Laminar Structure of the Porcine Hippocampus

**DOI:** 10.1101/201285

**Authors:** Alexandra V. Ulyanova, Paul F. Koch, Carlo Cottone, Michael R. Grovola, Christopher D. Adam, Kevin D. Browne, Maura T. Weber, Robin J. Russo, Kimberly G. Gagnon, Douglas H. Smith, H. Isaac Chen, Victoria E. Johnson, D. Kacy Cullen, John A. Wolf

## Abstract

The hippocampus is integral to working and episodic memory, and is a central region of interest in diseases affecting these processes. Pig models are widely used in translational research, and may provide an excellent bridge between rodents and non-human primates for CNS disease models due to their gyrencephalic neuroanatomy and significant white matter composition. However, the laminar structure of the pig hippocampus has not been well characterized. Therefore, we histologically characterized the dorsal hippocampus of Yucatan miniature pigs and quantified the cytoarchitecture of the hippocampal layers. We then utilized stereotaxis combined with single unit electrophysiological mapping to precisely place multichannel laminar silicon probes into the dorsal hippocampus without the need for image guidance. We used *in vivo* electrophysiological recordings of simultaneous laminar field potentials and single unit activity in multiple layers of the dorsal hippocampus to physiologically identify and quantify these layers under anesthesia. Consistent with previous reports, we found the porcine hippocampus to have the expected archicortical laminar structure with some anatomical and histological features comparable to the rodent and others to the primate hippocampus. Importantly, we found these distinct features to be reflected in the laminar electrophysiology. This characterization, as well as our electrophysiology-based methodology targeting the porcine hippocampal lamina combined with high channel count silicon probes will allow for analysis of spike-field interactions during normal and disease states in both anesthetized and future awake behaving neurophysiology in this large animal.

**Significance Statement:** The hippocampus is central to working and episodic memory and is critically affected by diverse disease processes. In order to investigate hippocampal electrophysiology in translational large animal models, we developed an imaging-free stereotaxis and intraoperative electrophysiology methodology with custom silicon probes to precisely localize probe placement within the hippocampal laminar structure. We report for the first time the profile of single units and local field potentials in the pig dorsal hippocampus and relate them to a histological description. This characterization forms the basis for accessible translational pig models to study diseases of the central nervous system affecting hippocampal circuitry in the large animal gyrencephalic brain, as well as the groundwork for potential awake behaving neurophysiology of the porcine hippocampus.

**Funding Sources:** The Department of Veterans Affairs, IK2-RX001479, I01-RX001097. The National Institutes of Health, NINDS R01-NS-101108-01, T32-NS043126. CURE Foundation, Taking Flight Award. DoD ERP CDMRP, W81XWH-16-1-0675.

## Introduction

The foundations of experimental neuroscience are largely rodent and non-human primate neuroanatomy, neurophysiology and neurochemistry. Rodents and non-human primates have also been the dominant species used to study neurological deficits involving the hippocampus such as memory disruption following traumatic brain injury (TBI), neurodegeneration and epilepsy (Buzsáki, 2015; Jang and Chung, 2016; Levin, 2003). While use of non-human primates reduces the problems due to species differences, these models are somewhat limited by economic and ethical restraints (Goodman and Check, 2002; Vink, 2017). Since gyrencephalic brain structure and appropriate grey/white matter ratios may be important for accurate modeling of CNS disorders, pigs have recently been proposed as an additional model for translational neuroscience research (Lind et al., 2007).

Much of the basic neuroanatomical characterization of the porcine brain and the hippocampus in particular has been done using both histological and clinical imaging methods such as positron emission tomography and magnetic resonance imaging (Lind et al., 2007; Rogers et al., 2008). These methods identified features from both the rodent and the non-human primate hippocampus such as its position in the brain, a laminated hilus and a dispersion of the deep pyramidal cells into stratum oriens similar to those of non-human primates (Holm and West, 1994; Lind et al., 2007; Sorensen et al., 2011). Recently, a comprehensive and detailed characterization of cortical surface anatomy and cytoarchitecture of the miniature pig’s brain was done based on Nissl-staining (Bjarkam et al., 2017). Here we histologically examine potential strain and age-related differences as well as the neuroanatomy of the miniature swine hippocampus in the sagittal and coronal plane.

Previous experimental and preclinical functional studies in both normal and pathophysiological states have been undertaken using functional MRI, electroencephalography and most recently both *in vivo* and *ex vivo* electrophysiology (Van Gompel et al., 2011; Wolf, 2017). As the value of the pig model in neurological diseases becomes more evident, further characterization of the anatomy and connectivity of the porcine hippocampus is required. In order to perform *in vivo* electrophysiological studies utilizing precisely placed high-density depth electrodes capable of recording laminar activity for comparative examination, we required a method of large animal stereotaxis without image guidance. We therefore implemented stereotaxis without the necessity of imaging to precisely place depth probes into the dorsal hippocampus of miniature pigs relative to the laminar structure of the CA1 region. We describe for the first time some of the basic characteristics of the electrophysiological laminar structure of the porcine hippocampus under anesthesia using custom high-density silicon laminar probes, including simultaneous recordings of single unit activity and local field potentials. We confirmed the laminar structure of the hippocampus and the precise location of the multichannel silicon probe using both electrophysiology and histopathology, reducing the need to immediately recover electrode tracks and increasing localization confidence during chronic implantations in future animals. Along with acute investigations, this methodology will ultimately allow for the potential expansion of hippocampal neurophysiology into awake behaving pigs under behaving conditions, as well as in chronic models of pathophysiological conditions such as traumatic brain injury and epilepsy.

## Materials and Methods

### Animals and Animal Care

In the current study, we used a total of 17 male Yucatan miniature pigs, which are known for their docile temperament, ease of handling, and slow growth rate (NSRRC Cat# 0012, RRID: NSRRC_0012). 2 of them were used for histological studies only, while 15 underwent both electrophysiological and histological examinations. Pigs were purchased from Sinclair and underwent the current studies at the approximate age of 5 - 6 months at a mean weight of 37.94 ± 3.26 kg (n = 17, mean ± SEM). We chose this age as the pigs are post-adolescent with near-fully developed brains, but young enough to be of a manageable weight for procedures and behavior (Duhaime et al., 2000; Flynn, 1984; Pampiglione, 1971). All pigs were pair housed when possible, but always in a shared room with other pigs. All animal procedures were performed in accordance with the [Author University] animal care committee’s regulations.

### Novel Porcine Stereotaxis

Although stereotactic approaches based on skull landmarks, or on individual subject imaging, have been designed for *in vivo* electrophysiological studies in pigs, they are either inconsistent, expensive to utilize, or difficult to implement (Felix, 1999; Marcilloux et al., 1989; Saito et al., 1998). Our goal was to design a relatively inexpensive stereotactic system combining a freely-available MRI atlas of pig anatomy (Saikali et al., 2010) with electrophysiological localization without the need for individual subject imaging in order to accomplish *in vivo* electrophysiological recordings across the laminar structure of the CA1 region of the pig hippocampus. Based on previous stereotactic work in our laboratory we chose to select bregma as the starting landmark for our stereotaxis. Bregma is defined as an anatomical point on the skull at which the coronal sutures intersect with the sagittal suture at the midline (Figure 2C). Although the relationship between bregma and the underlying brain can vary by several millimeters anterior to posterior (i.e. along the sagittal plane), the principle advantage is that bregma can be easily identified during surgery. We therefore defined the two-dimensional medial-lateral (ML) and anterior-posterior (AP) zero point to be bregma. Based on our previous experience we identified the approximate level of bregma in the coronal and sagittal planes on a freely available three-dimensional MRI atlas of the pig brain (Saikali et al., 2010). Figure 2A shows the orientation of the cortex and deeper structures, such as the dorsal and ventral hippocampus and the thalamus, at this level on the MRI atlas (also see Figure 1B). Based on this MRI atlas, we then chose 7 mm lateral to the midline as the sagittal plane that maximized CA1 thickness and aligned the dorsal hippocampal laminar structure perpendicularly with the cortical surface. Note that the Yorkshire (another pig strain) MRI atlas coronal section is structurally similar to a representative coronal section of the Yucatan brain used in this study, with slight shape variations due to strain differences (Figures 1B and 2A, see below).

**Figure 1.**
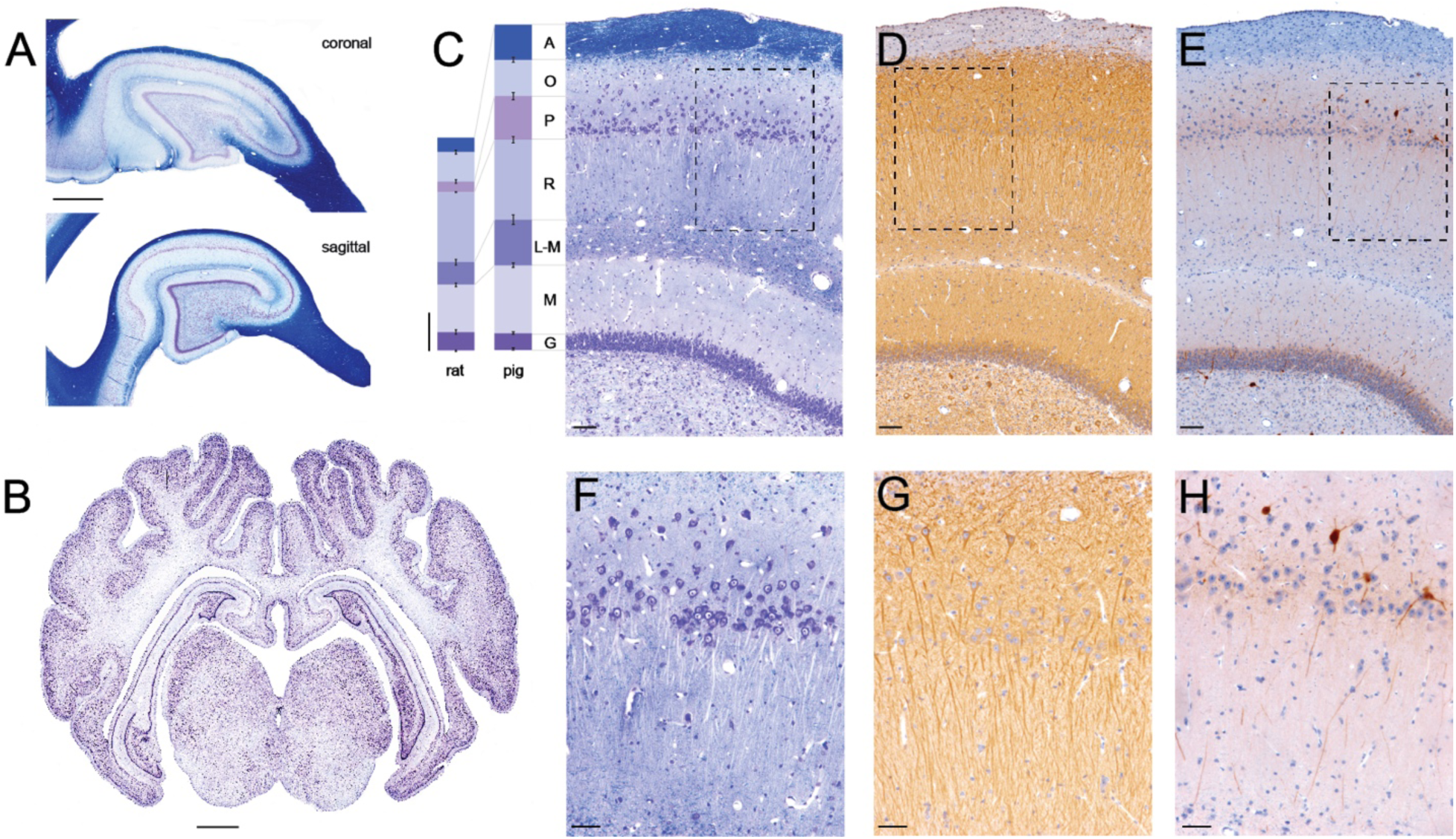
Morphology of Porcine Hippocampus. Sagittal and coronal sections of porcine brain were stained histologically to show organizational structure. **A)** Coronal (top) and sagittal (bottom) sections stained with LFB/CV show white and gray matter composition in dorsal hippocampus, which can be accessed stereologically from the brain surface. Both coronal and sagittal cuts through hippocampus produce arrow-like structures with consistent anatomical layers (scale = 2 mm). **B)** Coronal section stained with CV showing gyrencephalic structure (Bregma -1.5mm). Deep brain structures including hippocampus (dorsal and ventral) and thalamus are visible (scale bar = 2 mm). **C - E)** Hippocampal layers including the alveus (A), stratum oriens (O), stratum pyramidale (P), stratum radiatum (R), stratum lacunosum-moleculare (L-M), stratum moleculare (M), and stratum granulosum (G) are clearly visible (scale bar = 100 μm). **C)** Staining with LFB shows the myelinated axons, while counterstaining with CV highlights neurons in the pyramidal and dentate cell layers. Columns on the left show the depth of corresponding hippocampal layers quantified from H & E stained sections in Yucatan pigs (Bregma -1.5 mm; n = 8) and Long-Evans rats (Bregma -4.2 mm; n = 6). The depths for the pig hippocampal layers are (in μm, mean ± SEM,): A = 173.8 ± 11.55, O = 178.3 ± 15.01, P = 213.2 ±15.9, R = 394.1 ± 22.36, L-M = 223.1 ± 11.7, M = 335.1 ± 17.85, G = 81.3 ± 4.5. The corresponding depths for rat hippocampal layers are (in μm): A = 69.1 ± 9.05, O = 144.4 ± 7.51, P = 53.1 ±1.9, R = 343.9 ± 12.33, L-M = 112.2 ± 8.21, M = 232.8 ± 11.35, G = 90.6 ± 3.52 (scale bar = 200 μm). The following hippocampal layers were significantly larger in pigs than in rats: stratum alveus by 152 % (P < 0.001), pyramidale by 301 % (P < 0.001), radiatum by 15 % (P < 0.05), lacunosum-moleculare by 99 % (P < 0.001), and stratum moleculare by 44 % (P < 0.01), whereas the stratum oriens and granulosum were not significantly different. **D)** Staining with MAP2 identifies location of dendritic harbors, which are densely packed. **E)** A sub-population of interneurons is identified using parvalbumin (PV) label IHC in pyramidal and dentate granule cell layers. **F - H)** The pyramidal CA1 layer (scale bar = 50 μm). **F)** The pyramidal CA1 cell layer is widely dispersed, similar to the human CA1 layer, as shown with LFB/CV staining. **G)** Pyramidal CA1 cells send their dendrites down towards the hippocampal fissure as highlighted with MAP2 staining. **H)** PV^+^ interneurons located above, inside and below pyramidal CA1 layer, send their projections into stratum radiatum.

**Figure 2.**
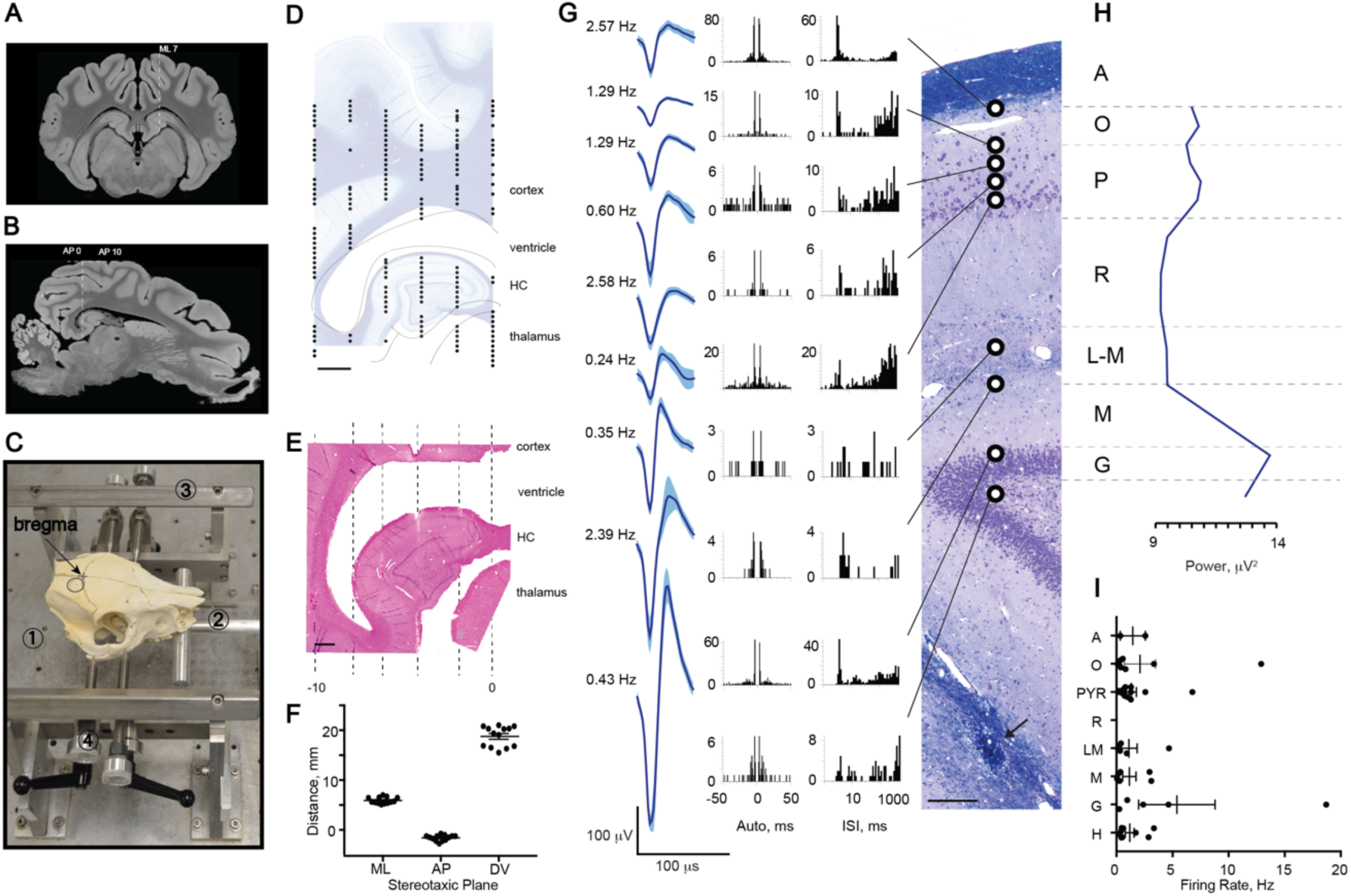
Single Unit Recordings in Porcine Hippocampus Corroborate Laminar Structure. **A)**. A representative coronal section from the MRI atlas (Bregma -1.5 mm) within the 12 mm AP range of interest shows the orientation of deep structures such as dorsal and ventral hippocampi (Saikali et al., 2010). We selected ML 7 as an access plane to the widest part of dorsal hippocampus. **B)** A sagittal section through ML 7 taken from the MRI atlas shows position and approximate size of the dorsal hippocampus. The image was rotated so that cortex above hippocampus is in the horizontal plane (horizontal dashed line). An approximate size of the hippocampus is shown between vertical dashed lines from right (AP 0) to left (AP -10). **C)** Photograph of a pig skull *in situ* in the novel stereotaxic instrument with an arrow pointing to Bregma and a circle indicating a craniectomy site. The main parts of the stereotaxis are frame base (1), bite bar (2), anterior-posterior (AP) bars (3), and pins (4). The animal’s head is first guided between AP bars and onto an adjustable bite bar, then a set of four pins is inserted into the zygomatic bones. Location and size of the craniectomy are shown (black circle). **D)** ML coordinates obtained with this stereotax are fairly accurate, but AP coordinates are not. Therefore, electrophysiological mapping is needed for each experiment. The digitized tungsten map from our initial animal is shown in the AP plane at ML 7 overlying a representative sagittal section, stained with LFB/CV (scale bar = 2 mm). The lines outline deep brain structures including subcortical white matter, ventricle, hippocampus and thalamus. **E)** A representative sagittal section from the initial animal stained with H&E (scale bar = 1 mm) contains Hamilton syringe tracks in the AP plane in 2 mm steps from AP 0 to AP -10 (dashed lines). **F**) The precision and reproducibility of our targeting technique in 13 animals was estimated histologically in the ML and AP planes and electrophysiologically in the DV plane. The estimated distances are (in mm): ML = 5.91 ± 0.17 from midline, AP = -1.60 ± 0.16 from bregma, and DV = 18.76 ± 0.57 (mean ± SEM, n=13) from brain surface to first multi-unit activity. **G - I)** Example of single spiking activity during the mapping procedure performed by slowly advancing a Tungsten electrode along the indicated track. **G)** A coronal section of the dorsal hippocampus stained with LFB/CV is shown from a subsequent animal from which Tungsten electrode recordings were made during the initial mapping procedure (scale bar = 100 μm). Arrow points to the subsequently inserted silicon probe artifact. Histologically identified layers are labeled: alveus (A), stratum oriens (O), stratum pyramidale (P), stratum radiatum (R), stratum lacunosum-moleculare (L-M), stratum moleculare (M), stratum granulosum (G). Three minute recordings were made at regular intervals spaced 100 - 200 μm apart. Raw neural signals were processed offline and putative single units were isolated. Depths at which single units were successfully isolated are indicated by black and white circles. Representative sing units from each hippocampal layer are shown as averaged waveforms (mean ± SD) accompanied by firing rates, autocorrelograms and inter spike interval histograms (Robertson et al.). **H)** To capture all multi-unit activity and confirm the locations of the cellular layers, the RMS power of the high frequency bandpass filtered signal (600-6,000 Hz) was calculated. A representative profile of spiking activity shows several peaks at the corresponding pyramidal and granular cell layers. **I)** Firing rates of single units recorded in 2 animals with Tungsten electrodes were averaged within each of 8 layers spanning approximately 200 μm each, corresponding to the hippocampal layers calculated in Figure 1C (mean ± SEM, n = 2).

Because the position of bregma in the AP plane with respect to the brain varies between animals, we selected an approximate range of AP coordinates that spans the AP extent of the dorsal hippocampus on the MRI atlas, approximately 12 mm in total. A sagittal section through ML 7 taken from the MRI atlas shows the position and approximate size of the dorsal hippocampus (Figure 2B, horizontal line, see below). The image is rotated so that cortical surface above the hippocampus is horizontal. Again, the sagittal section obtained from the MRI atlas is similar to the sagittal histological section of porcine brain used in our study (Figure 1A, bottom). This section of brain defined by the AP extent of the dorsal hippocampus at ML 7 then became the region of interest for electrophysiological mapping during surgery (see below).

### Placement in Stereotactic Frame

We constructed a novel stereotactic frame based on percutaneously placed skull pin fixation, a commonly employed technique in human neurosurgery (Figure 2C, see below). The stereotactic frame consisted of four major components: frame base, bite plate, anterior-posterior (AP) bars, and skull pins (Figure 2C, see below). The frame base (1) hold all parts of the stereotax and was made of heavy aluminum alloy with holes to attach the AP bars of the stereotactic frame. As the animal was guided into the stereotactic frame, its jaw was opened and its maxilla placed on top of the bite bar (2) with the endotracheal tube positioned below the bar. The point at which the bite bar meets the frame base was adjustable. Additionally, the bite bar could be maneuvered in the AP plane and rotated in the coronal plane, allowing the researcher to optimally position the head prior to pin insertion. The AP bars (3) were independent parallel rails fastened to the frame base and run parallel to the animal’s head in the sagittal plane. These arms had standard 1 mm markings. Stereotactic manipulator arms (Kopf Instruments, Cat# 1760) could then be secured to the AP bars and moved along them in the AP plane using the Vernier scale to 0.1 mm precision. Before securing the animal with the skull pins, two points on each side of the animal along the zygoma, approximately 3 cm apart, were sterilely prepped with chlorhexidine solution and a small stab incision of 2-3 mm made at each point with a scalpel. Four stainless steel pins (4) affixed to the frame were inserted into the zygoma at each of these points providing rigid fixation of the head to the frame. In order to position the head precisely for stereotactic localization, each of these pins could be moved in three dimensions while the pig was secured in the stereotax.

We used an iterative process of progressively finer adjustments both before and after skull pin fixation to place the pig’s head precisely within the frame at a consistent angle such that the dorsal surface of the brain above the hippocampus was parallel to the base of the stereotactic frame. The pig was first positioned approximately within the frame using a bite bar. The four skull pins were then inserted through the skin into the bone of bilateral zygomas, fixing the head. The head was then further adjusted to the desired angle using the skull pins, which can be moved in three dimensions. A final fine adjustment to the skull angle was made via direct measurements on the surface of the skull using a stereotactic arm and measurements on the frame once the skull had been exposed.

An angle of 10–12° between the skin above the dorsal surface of the skull and the base of the stereotax was initially selected so that the cortical surface above the hippocampal region is level with the AP bars (Figure 2B, dashed line, see below). Following initial alignment and skull fixation of the animal in the stereotactic frame to achieve this angle, the animal’s head was re-positioned again after skin incision based on fine measurements of the skull surface and bregma. An electrode holder with a marking pin was assembled and zeroed to bregma and the skull surface. In order to level the skull in the coronal plane, the dorsal-ventral (DV) coordinates were first measured in the medial-lateral (ML) plane. Head angle was finely adjusted if the difference in the DV coordinates on each side of the skull at ML = 10 mm was more than 0.5 mm. Next, the DV coordinates were measured in the midline at bregma and AP = +30 mm. The angle between the skull surface and the base of stereotax was finely adjusted by repositioning the pins until the difference in the DV measurements at these two points was no more than 2.9 ± 0.1 mm, which corresponds to an angle of 7°. We found this angle to be ideal for reproducibly locating and mapping the dorsal hippocampus.

### Surgical Procedure

Yucatan miniature pigs were fasted for 16 hours then induced with 20 mg/kg of ketamine and 0.5 mg/kg of midazolam. Animals were intubated with an endotracheal tube and anesthesia was maintained with 2-2.5% isoflurane per 2 liters O_2_. Each animal was placed on a ventilator and supplied oxygen at a tidal volume of 10 mL/kg. A catheter was placed in an auricular vein to deliver 0.9% normal saline at 200 mL per hour. Additionally, heart rate, respiratory rate, arterial oxygen saturation, end tidal CO_2_ and rectal temperature were continuously monitored, while pain response to pinch was periodically assessed. All of these measures were used to titrate ventilation settings and isoflurane percentage to maintain an adequate level of anesthesia. A forced air warming system was used to maintain normothermia throughout the procedure.

15 pigs in this study underwent electrophysiological recordings and analysis. These pigs were placed in a stereotactic frame (described above), with the surgical field prepped and draped. A linear incision was made along the midline. A 13-mm diameter burr hole was made centered at 7 mm lateral to the midline and 4.5 mm posterior to bregma. The bony opening was subsequently expanded using Kerrison punches. The dura was opened in a cruciate manner using a #11 blade.

### Electrophysiological Mapping of Porcine Hippocampus

In all 15 animals, the dorsal hippocampus was mapped with several Tungsten electrode insertions in the sagittal plane, utilizing observed spiking activity to generate a two-dimensional map (Figure 2D, see below). For this mapping, we used a high impedance Tungsten monopolar electrode (Ø = 125 μm, length = 60 mm, impedance = 0.5 MΩ; FHC, Cat# UEWSEGSEBNNM). A skull screw was placed over the contralateral cortex as a reference signal.

We then chose coordinates within this map for insertion of either a custom-designed linear 32-channel silicon Edge probe with 200 μm spacing (NeuroNexus, Cat# V1x32-Edge-10mm-200-312-Ref; n = 4) or a custom-designed 32-channel silicon Vector probe (NeuroNexus, Cat# V1x32-15mm-tet-lin-177; n = 11) such that the spread of electrode contacts would span the laminar structure of the dorsal hippocampus at its maximal thickness in the dorsal-ventral plane, perpendicular to the individual hippocampal layers. The silicon probes were designed with one low-impedance channel placed 2 mm above the next most-proximal channel, which we used as a reference signal (Figure 3A, see below). For dorsal hippocampal targeting, this usually results in the reference channel being positioned within the temporal horn of the lateral ventricle sitting just above the hippocampus and provides 31 channels for intra-hippocampal recordings. The custom-designed silicon Vector probe was similar to the custom-designed Edge linear silicon electrode, but the spatial arrangement of the 31 intra-hippocampal channels were of a custom design (data not shown). At the end of the procedure the anesthetic level was deepened (to 5% isoflurane) and pigs were sacrificed for histological analysis.

**Figure 3.**
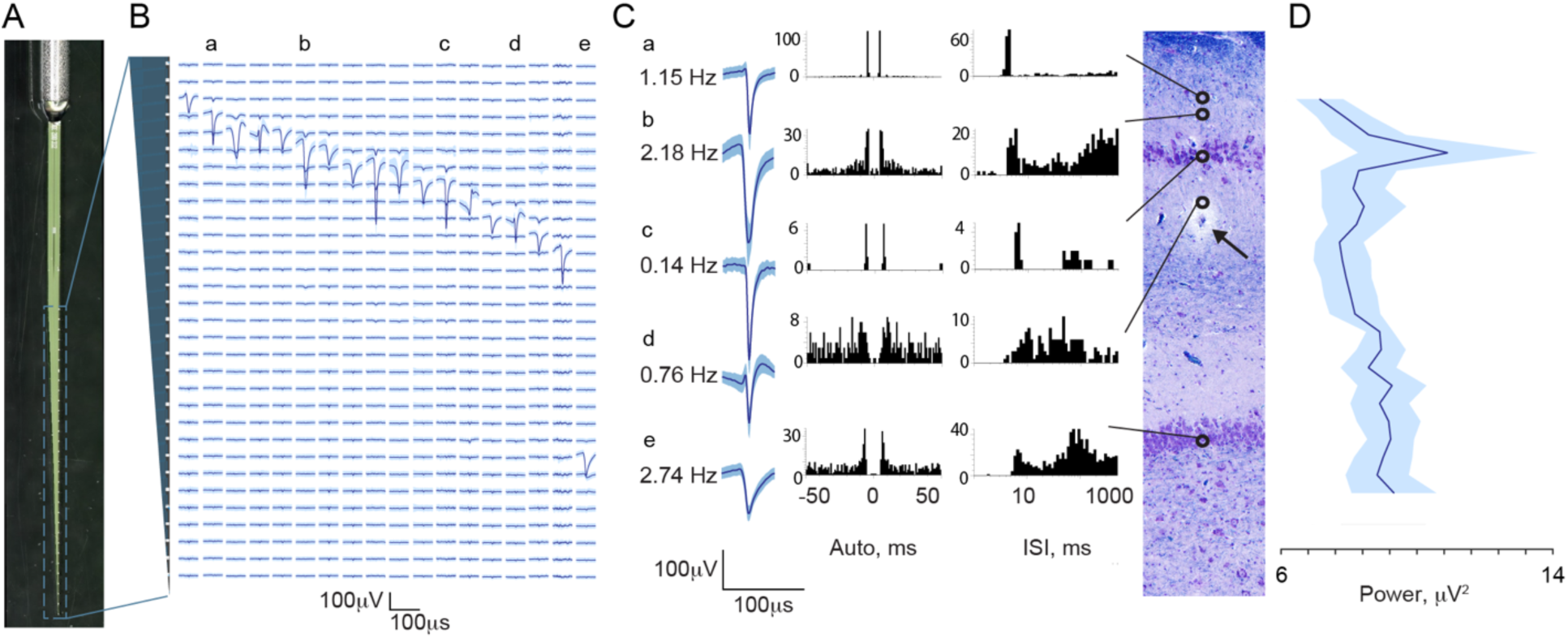
Single Units Recorded with Silicon Laminar Probes Confirm Laminar Structure. **A)** Photograph of the custom-made Edge silicon probe shows an internal reference and linear orientation of its 31 channels. In each animal, this probe was inserted into the dorsal hippocampus based on initial electrophysiological mapping. Electrical activity (B-D) is shown as recorded from the final position of the probe. **B)** Example in one animal of the laminar profile of single unit activity shown as averaged waveforms (mean ± SD) aligned to the 31 recording channels of the silicon probe along the vertical axis (200 μm spacing), indicated by the gray cartoon of the probe on the left. Lowercase labels (a-e) indicate single unit examples displayed in C. **C)** A coronal section of the dorsal hippocampus stained with LFB/CV is shown from the same animal (scale bar = 100 μm). Arrow points to silicon probe artifact, at approximately mid-shaft. Open circles indicate channels on which single spiking units were isolated revealing a large cluster in the region of stratum pyramidale and a single channel in the region of stratum granulosum (scale bar: X axis – 2 msec, Y axis - 100 μV). Examples of isolated spike displayed as averaged waveforms (mean ± SD) are shown along with autocorrelograms and interspike intervals histograms (Robertson et al.). Alignment of single unit examples to the histology was based on recovered artifact of the probe tip (not shown), stereotactic coordinates used during insertion and known spacing of the 31 contacts. Vertical alignment is only approximate as the plane of insertion was oblique to the histological coronal plane and the relative thicknesses of the hippocampal layers change along the AP axis. **D)** Stratum pyramidale of CA1 was identified in four animals recorded with the Edge laminar silicon probe based on the maximum power of multi-unit spiking activity (RMS power in the 600-6,000 Hz band) (mean ± SEM, n = 4). The traces from all four animals aligned at the peak of the RMS power are shown. Only aligned channels where all four animals overlapped are included (total channels displayed = 25 out of 31).

### Neural Data Collection and Analysis

All electrophysiological recordings were made under isoflurane anesthesia (2 - 2.5 %). Neural signals were amplified and acquired continuously at 32 kHz on a 64-channel Digital Lynx 4SX acquisition system with Cheetah recording and acquisition software (Neuralynx, Inc.).

#### Spike Detection and Analysis

Signals acquired from the tungsten monopolar electrode during the mapping procedure were bandpass filtered (600 Hz to 6 kHz) and thresholded for spike detection in real time. Thresholds were chosen based on observed signal to noise ratios during the session. Recorded spike trains and waveforms then underwent off-line automated spike sorting using KlustaKwik software http://klusta-team.github.io/klustakwik/, RRID: SCR_014480) and further manual refinements using SpikeSort3D (Neuralynx, Inc.). Neural signals acquired from the 31 channels on the silicon probe were bandpass filtered (0.1 Hz to 9 kHz) in real time prior to sampling. Off-line spike detection and sorting was performed on the wideband signals using the Klusta package (http://klusta-team.github.io/klustakwik/, RRID: SCR_014480), which was developed for higher density electrodes, and manually refined with KlustaViewa software (https://github.com/klusta-team/klustaviewa). The Klusta routines are designed to take advantage of the spatial arrangement of and spike timing from all probe channels simultaneously in constructing putative clusters (Rossant et al., 2016). Resulting single-unit clusters were then imported into Matlab software, version R2017a for visualization and further analysis using custom and built-in routines (https://www.mathworks.com/products/matlab.html, RRID: SCR_001622). In order to minimize waveform shape distortion for visualization, the wideband signal was high pass filtered using a wavelet multi-level decomposition and reconstruction filter (level 6, Daubechies 4 wavelet) (Wiltschko et al., 2008). Waveforms were then extracted from this filtered signal and averaged for display. Spike trains were imported into NeuroExplorer software, version 6 for further analysis and to generate autocorrelograms and interspike intervals histograms (http://www.neuroexplorer.com/, RRID:SCR_001818).

#### Analysis of Local Field Potentials (LFP)

Acquired wideband LFP recorded from 31 channels of the silicon probe were down-sampled to either 2 or 3 kHz for further analysis. Signals were imported into Matlab software, version R2017a (https://www.mathworks.com/products/matlab.html, RRID: SCR_001622) and processed using a combination of custom and modified routines from the freely available Matlab packages FMAToolbox (http://fmatoolbox.sourceforge.net, RRID:SCR_015533), Chronux (http://chronux.org, RRID: SCR_005547), and EEGLAB (http://sccn.ucsd.edu/eeglab/index.html, RRID:SCR_007292) (Hazan et al., 2006; Mitra, 2007). Current source density (CSD) was calculated as the second spatial derivative of the LFP across all 31 hippocampal channels and visualized by convention as a heat map of current sinks (blue) and sources (orange) (Mitzdorf, 1985) using the freely available Matlab package CSDplotter, version 0.1.1 (Pettersen et al., 2006).

To perform spectral power analysis across laminar structure, the hippocampal cellular layers were first identified in four animals recorded with the Edge laminar probe by calculating the root mean squared (RMS) power in the 600-6,000 Hz frequency band. The traces from multiple animals (n = 4) were then aligned at the peak of 600-6,000 Hz band, which corresponds to the pyramidal CA1 cell layer and non-overlapping channels were cut off (total channels displayed = 25, out of 31, n = 4). After alignment, recordings from the 25 remaining hippocampal channels were broken into 1 sec segments. Each segment underwent multi-taper spectral analysis in the frequency ranges of theta (4 - 10 Hz), low gamma (25 - 55 Hz), high gamma (55 - 90 Hz), and a ripple frequency range (150 - 250 Hz) at each channel (Chernyy et al., 2008). The power for each 1 sec segment at each channel was then integrated over the entire frequency range. The values for each channel were then averaged across all 1 second segments resulting in the average integrated spectral power as a function of probe depth. For each animal, power spectral density (PSD) analysis of the channel presenting the maximum deflection during sharp-wave-like events was calculated in 1-20 Hz frequency band using the Welch’s method with the following parameters: window length = 1 sec, frequency resolution = 0.2 Hz, overlap = 60%. Each PSD was then normalized by its peak value and the average displayed as mean ± SEM.

### Tissue Handling and Histological Examinations

Histological analyses were performed on brain tissue from male Yucatan miniature pigs (n = 17 total) in order to locate tracks generated by electrode insertion (n = 15), or to histologically characterize the porcine hippocampus (n = 2). While under 5 % isoflurane anesthesia, all animals underwent transcardial perfusion with 0.9% heparinized saline followed by 10% neutral buffered formalin (NBF). After post-fixation for 7 days in 10% NBF at 4°C, each brain (weight = 88.29 ± 2.12 g, mean ± SEM, n = 17) was dissected into 5 mm blocks in the coronal (n = 15) or sagittal (n = 2) planes and processed to paraffin using standard techniques. 8 μm sections were obtained at the level of the hippocampus in either the coronal or sagittal planes. Standard hematoxylin & eosin (H&E) staining was performed on all animals (n = 17) to visualize electrode tracks. Since electrode insertion could potentially disturb the cytoarchitecture, we utilized the hemisphere contralateral to the electrode insertion for histological estimation of hippocampal layers.

#### Luxol Fast Blue / Cresyl Violet (LFB/CV) Staining

Tissue sections were dewaxed in xylenes and rehydrated to water via graded ethanols before being immersed in 1% LFB solution (Sigma, S3382) at 60°C for 4 hours. Excess stain was then removed by immersion of sections in 95% ethanol. Differentiation was performed via immersion in 0.035% lithium carbonate for 10 seconds followed by multiple immersions in 70% ethanol until the gray and white matter could be clearly distinguished. Slides were rinsed and counterstained via immersion in preheated 0.1% CV solution (Sigma, C5042) for 5 minutes at 60°C. After further rinsing, slides were differentiated in 95% ethanol with 0.001% acetic acid, followed by dehydration, clearing in xylenes and cover slipping using cytoseal-60.

#### Immunohistochemistry (IHC)

Single IHC labeling was performed according to previously published protocols. Briefly, tissue sections were dewaxed in xylenes, rehydrated to water via graded ethanols and immersed in 3% aqueous hydrogen peroxide for 15 minutes to quench endogenous peroxidase activity. Antigen retrieval was achieved via microwave pressure cooker at high power for 8 minutes, submerged in Tris EDTA buffer (pH 8.0). Sections were then incubated overnight at 4°C using antibodies specific for MAP2 (Abcam, Cat# ab5392 Lot# RRID: AB_2138153, 1:1000), parvalbumin (Millipore Cat# MAB1572 Lot# RRID: AB_2174013, 1:1000), and the N-terminal amino acids 66–81 of the amyloid precursor protein (APP) (Millipore, Cat# MAB348 Lot# RRID: AB_94882, 1:80K). Slides were then rinsed and incubated in the relevant species-specific biotinylated universal secondary antibody for 30 minutes at room temperature. Next, application of an avidin biotin complex (Vector Laboratories, Cat# PK-6200 Lot# RRID: AB_2336826) was performed for 30 minutes, also at room temperature. Lastly, the 3, 3’-diaminobenzidine (DAB) peroxidase substrate kit (Vector Laboratories, Cat# SK-4100 Lot# RRID: AB_233638) was applied according to manufacturer’s instructions. All sections were counterstained with hematoxylin, dehydrated in graded ethanols, cleared in xylenes, and cover slipped using cytoseal-60.

### Slide Imaging

Whole brain coronal tissue sections were digitally scanned at 20X magnification using the Aperio CS2 Digital Pathology Scanner (Leica Biosystems). Additional images were captured on a Nikon Eclipse 80i upright microscope and a DSRi1 camera (Nikon, Inc.), with associated NIS-Elements software, version V4.5 (https://www.nikoninstruments.com/Products/Software, RRID: SCR_014329).

### Comparative Analysis of Hippocampal Architecture

Comparative analysis of hippocampal layers between species was done on H&E stained sections. Hippocampal layers were measure on either scanned coronal sections of porcine brain (n = 8) or imaged coronal sections of archived formalin-fixed paraffin-embedded rat brain from male Long-Evans rats (n = 6). Whole brain coronal sections at hippocampal level were stained with H&E and imaged as described above. Individual hippocampal layers were visually identified and measured.

### Statistical Analysis

The data was analyzed using Graphpad Prism software, version 7 (http://www.graphpad.com/, PRID: SCR_002798). The histological analysis of hippocampal cytoarchitecture in miniature Yucatan pigs (n = 8) and Long-Evans rats (n = 6) is presented as mean ± SEM. For comparative analysis of hippocampal layers, corresponding depths of hippocampal layers in pig were compared to those in rats (t-test). The electrode placement and stereotactic precision is presented as mean ± SEM (n = 13). Individual single unit waveforms recorded with both Tungsten electrodes and silicon laminar probes are presented as mean ± SD. Analysis of single unit firing rates recorded with Tungsten electrodes were averaged over 200 μm distance and are presented as mean ± SEM, n = 2 pigs. Power spectrum analysis was performed in animals implanted with linear Edge probe and is presented as averaged power spectrum of various frequency bands (mean ± SEM, n = 4).

## Results

### Neuroanatomy of Porcine Hippocampus

In order to plan our initial stereotactic approach to the dorsal hippocampus in the Yucatan miniature pig, we first undertook a gross anatomical and histological study of the structure. Three Yucatan miniature pigs were sacrificed and the brains extracted (see Methods). Figure 1B shows the sulci, multiple gyri and white/gray matter composition of the porcine brain in coronal section (Bregma -1.5 mm) stained with CV, highlighting subcortical structures such as the hippocampus and thalamus that are stereotactically accessible from the brain surface. Analogous to rodent and human hippocampus, porcine hippocampus is oriented so that both coronal and sagittal planes produce arrow-like structures from anatomical layers in a similar orientation, as shown with LFB staining in Figure 1A. The granule cells of the dentate gyrus form the point of the arrow, the notably large hilar region in pigs forms the body of the point and the pyramidal cells of the cornu ammonis (CA) regions form the curved shaft in both planes. The classic hippocampal layers (alveus (A), stratum oriens (O), stratum pyramidale (P), stratum radiatum (R), stratum lacunosum-moleculare (L-M), stratum moleculare (M), and stratum granulosum (G)), as well as the hippocampal fissure, which divides the dentate from the CA regions, are clearly visible under higher magnification (Figure 1C-E).

The width of each hippocampal layer was quantified from histologically stained sections of pig brain tissue used in this study (Bregma -1.5 mm) and compared to archived, histologically stained sections of rat brain tissue at the similar hippocampal level (Bregma -4.20 mm) (see Figure 1C). Columns show the widths of corresponding hippocampal layers quantified from H & E stained sections in pigs (mean ± SEM, n = 8) and rats (mean ± SEM, n = 6). The corresponding widths for pig hippocampal layers are (in μm): A = 173.8 ± 11.55, O = 178.3 ± 15.01, P = 213.2 ±15.9, R = 394.1 ± 22.36, L-M = 223.1 ± 11.7, M = 335.1 ± 17.85, G = 81.3 ± 4.5. The corresponding widths for rat hippocampal layers are (in μm): A = 69.1 ± 9.05, O = 144.4 ± 7.52, P = 53.12 ±1.9, R = 343.9 ± 12.33, L-M = 112.2 ± 8.21, M = 232.8 ± 11.35, G = 90.6 ± 3.52. The following hippocampal layers were significantly larger in pigs than in rats: stratum alveus by 152% (P < 0.001), pyramidale by 301 % (P < 0.001), radiatum by 15 % (P < 0.05), lacunosum-moleculare by 99 % (P < 0.001), and moleculare by 44 % (P < 0.01), whereas hippocampal layers stratum oriens and granulosum were not significantly different.

Staining with LFB/CV highlights myelinated fibers and neuronal cell bodies in the pyramidal and granular cell layers, revealing the expected ordered orientation of these fibers in relation to the cell bodies, a feature that plays critically in the laminar interpretation of hippocampal electrophysiology (Figures 1C, F). The pyramidal CA1 cell layer appears less densely packed in pigs due to a progressive dispersion of these cells into stratum oriens in the direction of the subiculum as previously reported (Holm and West, 1994). This dispersion has the effect of partially blurring the distinction between stratum pyramidale and the most dorsal aspects of stratum radiatum, as synaptic inputs into the dendritic tree of the more dorsally-located cells will co-localize with the more ventrally-located cell bodies. This phenomenon can be readily seen with MAP2 labelling, showing pyramidal cells above the main CA1 layer sending dendrites through stratum pyramidale into the distal layers (Figure 1D, G). These dendritic arbors are very dense and intermingled. Finally, as previously reported in another strain of pig, a sub-population of interneurons located in the pyramidal and dentate granular cell layers can be identified with Parvalbumin (PV^+^) staining (Figure 1E, H) (Holm et al., 1990). These PV^+^ positive interneurons primarily located above, inside, and below the pyramidal CA1 layer also have deep projections into the radiatum.

### Targeting the Dorsal Hippocampus Using Electrophysiological Mapping

We use electrophysiological mapping to address two different scales of localization: the localization of the probe within the three-dimensional structure of the hippocampus, largely in the medial-lateral and anterior-posterior planes, and the laminar localization of the probe with respect to the dendritic inputs along the dorsal-ventral axis of CA1. Since the initial, atlas-based coordinates selected for electrode placement can be inaccurate due to skull and individual animal variation as well as minor brain swelling during surgery, the precise location of the hippocampus needs to be determined within the stereotactic space. We utilized an electrophysiological mapping technique similar to that routinely employed clinically during deep brain stimulation implantation procedures (Gross et al., 2006), where a functional map of spiking activity in the cortex, dorsal hippocampus and top of the thalamus was created by driving a small diameter (Ø = 125 μm) high impedance Tungsten electrode into the brain and recording the coordinates at which neuronal spiking activity was detected.

We created a functional map of cortex and hippocampus by recording neuronal spiking activity at various depths, and we confirmed that electrophysiological activity of the dorsal hippocampus matches the anatomy using histological needle artifacts from stereotaxic dye injections (Figure 2). In our first experiment, mapping in our initial animal was executed in the sagittal plane 7 mm lateral to midline by advancing a single Tungsten electrode in equal steps in the AP plane in 2 mm intervals (AP range: 0 – 10 mm). At each AP level, we drove the electrode from the brain surface (DV = 0) to a final depth of 30 mm (DV = 30). We notated where multi-unit spiking activity was detected along each track, thereby creating a two-dimensional map of the dorsal hippocampus and surrounding structures in the sagittal plane (Figure 2D, circles). The map of spiking activity generated an outline of the ventricle, dorsal hippocampus, thalamus and cortex, shown overlying a representative sagittal section stained with LFB/CV (Figure 2D). In order to assess whether the generated outline of electrophysiological activity corresponded to the histology, blue dye was stereotactically injected through a Hamilton syringe (26-gauge needle, outer diameter Ø = 0.46 mm) along the same tracks as the Tungsten electrode in the sagittal plane to a depth of 30 mm (n = 2). After sacrificing the animal, the brain was cut in the sagittal plane at the level of the hippocampus, revealing the blue dye tracks (not shown). We then cut serial sections at this level in the sagittal plane and stained them with H&E, locating the track artifacts of the injection needle (Figure 2E). We aligned the generated electrophysiological map with the sagittal section best showing the track artifacts (Figure 2E, dashed lines), confirming accurate placement within the dorsal hippocampus. In subsequent experiments the generated electrophysiological map was used to place a multichannel silicon probe within the dorsal hippocampus for subsequent experimental recordings (discussed below). In these animals, while electrophysiological mapping was still performed in the sagittal plane, brains were sectioned in the coronal plane and sections demonstrating artifact from the silicon probe were located (Figures 3 and 4, see below). After our initial experiments this technique became routine and we were able to use fewer tracks on later animals (as few as three) to localize the hippocampus in sagittal profile.

**Figure 4.**
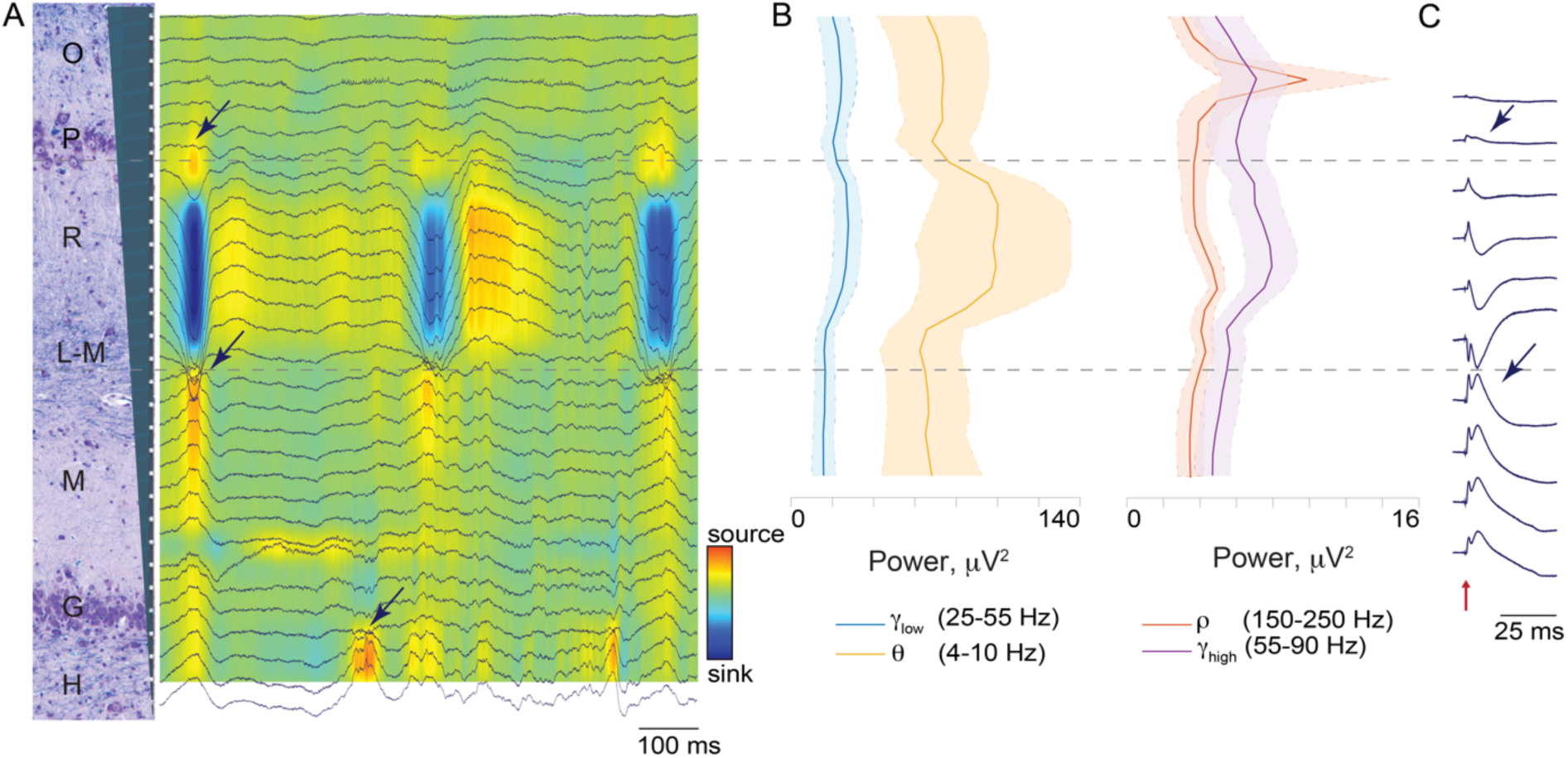
Laminar Structure of Extracellular Recordings Reveals Electrode Position. **A)** Tracing of wide-band local field potentials (LFP) across all channels aligned to the 31 recording positions of the silicon Edge probe along the vertical axis (200 μm spacing) represented by the gray cartoon on the left. The signal was referenced to the skull ground screw. Multi-unit spiking activity can be seen in a dorsal band indicating stratum pyramidale (approximately channel 4). Note regular bursts of synchronized activity that occur approximately every 300 ms, potentially representing sharp wave-like activity under isoflurane anesthesia (Lustig et al., 2015) (scale bar: X axis – 100 ms, Y axis - 200 μV). Arrows point to the phase reversal of the LFP above and below stratum radiatum as well as at dentate gyrus middle molecular layer (scale bar: X axis – 25 ms, Y axis - 200 μV). CSD map shows rhythmic large sink in stratum radiatum. **B**) Power spectra of classic theta (orange, 4-10 Hz), low gamma (blue, 25-55 Hz), high gamma (purple, 55-90 Hz), and the ripple range (red, 150-250 Hz) bands are shown aligned to the channels in A. Each power band is presented as mean ± SEM (n = 4). Note that the theta band (orange) has maximal power in stratum radiatum. Although we found maximal power in the ripple band in stratum pyramidale (by definition the channel of maximal multi-unit spiking activity; see Figure 3D), we did not observe any definitive, discrete ripple oscillatory events. **C)** Responses to perforant path stimulation (red arrow) were averaged over a 50 msec time period (20 stimulations, 2 seconds apart, stimulation artifact removed for clarity). The laminar profile of these evoked potentials is shown in one animal. Phase reversals were observed above and below stratum radiatum (arrows).

In order to assess the precision and reproducibility of our targeting technique within the three-dimensional structure of the dorsal hippocampus, we estimated the final position of the inserted silicon probe histologically in the ML and AP planes and electrophysiologically in the DV plane (Figure 2F, mean ± SEM, n=13). In the ML plane, the average measured histological distance from the point of insertion into the dorsal hippocampus to the midline (in mm) was 5.91 ± 0.17 (mean ± SEM, n = 13). Compared to our stereotactic ML insertion point of 7 mm from midline, this is a discrepancy of 16%, which is within the 11-25% shrinkage range reported in the literature after formalin fixation (Quester and Schröder, 1997). In the AP plane, we estimated the position of the point of insertion into the dorsal hippocampus by comparing the histological coronal section depicting the electrode tract with the MRI atlas (Saikali et al., 2010), choosing the coronal MRI slice (0.25 mm spacing) that best matched the overall hippocampal morphology in the histology. The average AP position of our probes in MRI space (in mm) was -1.60 ± 0.16 (mean ± SEM, n = 13). As a visual reference, the coronal MRI slice depicted in Figure 2A has an AP coordinate of -1.00. Unlike localization in the ML and AP planes, our determination of DV precision was made by the electrophysiology. During insertion of the silicon probe, the most dorsal point within the dorsal hippocampus where neuronal spiking activity was detected, corresponding either to neurons within stratum oriens, or to the pyramidal CA1 layer was located an average of 18.76 ± 0.57 mm (mean ± SEM, n = 13) below the brain surface in our stereotactic space.

### Hippocampal Laminar Structure and Intra-Hippocampal Electrode Localization in the Dorsal-Ventral Axis

Localizing within the AP and ML axes (in the sagittal and coronal planes, respectively) of the dorsal hippocampus as described above could also be accomplished with individual subject imaging and fiducial registration. However, localizing our probe precisely in the dorsal-ventral (DV) axis relative to the spiking activity in the CA1 pyramidal layer and dentate gyrus, as well as to the laminar structure of dendritic inputs between them, is more relevantly accomplished with electrophysiology, as neural activity is the direct measurement of interest. In two animals, once the electrophysiological map in the sagittal plane was created, we then recorded with the same high impedance electrode at successive depths along a single track of our map in order to isolate and characterize the spiking activity as a function of depth. We advanced the electrode along this track in 100 - 200 μm intervals, recording for 3 minutes at each level. We then isolated putative single units off-line. A larger multichannel silicon probe was then inserted to a depth of 25 mm along this same track as shown in Figure 2G. The stereotactic coordinates of this probe as well as of the generated map, along with the histological artifact generated by the silicon probe (Figure 2G, black arrow), allowed us to approximately align the map of recorded single unit activity with a representative coronal section from the same animal stained histologically with LFB/CV showing the probe artifact (coronal plane). This single electrode data revealed a cluster of single unit activity aligned with the diffuse pyramidal cell layer depicted in the histology and a second cluster of single unit activity aligned to the edge of the dentate gyrus at the granular cell layer and hilus. While various cell types in the hippocampus of the domestic pig have been identified with immunohistological markers previously, their electrophysiological characteristics are unknown (Holm et al., 1992; Holm et al., 1993; Holm et al., 1990). Here we report representative average waveforms (mean ± SD), autocorrelograms, inter spike intervals histograms as well as firing rates of putative single neuron action potentials within the various porcine hippocampal layers (Figure 2G). The locations of the cell body layers in this animal were also confirmed by calculating multi-unit spiking activity power in the 600-6,000 Hz frequency band (Figure 2H). This demonstrates peaks in stratum pyramidale (P) and stratum granulosum (G). The firing rates of the individual single units recorded in both animals were then averaged in spatial bins of approximately 200 μm in depth (Figure 2I), corresponding to the depths of the hippocampal layers calculated in Figure 1C (mean ± SEM, n = 2).

### Simultaneous Laminar Field Potentials and Single Unit Activity via High Density Silicon Probe

In a total of 15 animals, once the map of spiking activity of the dorsal hippocampus is created in the sagittal plane, we were then able to place a multichannel silicon probe spanning the lamina of CA1 based on this map. In these experiments, we chose a point in the AP plane where we determined both that the CA1 layers were most perpendicular to the inserted probe and that showed the thickest span of spiking activity in the pyramidal layer along the DV axis in our electrophysiological map. We then inserted either the custom Vector, or custom Edge 32 channel silicon probe (Edge probe depicted in Figure 3A: 31 recording sites, 200 μm spacing) into the dorsal hippocampus based on the initial mapping. We advanced the probe while monitoring spiking activity and the local field potential (LFP) on the multichannel probe in real time. In this way, we placed the large band of spiking activity corresponding to the pyramidal cell layer near the top of the probe (Figure 3B). The internal reference on the silicon probe, located 2 mm above the first channel, is placed in the ventricle. Once in position, we recorded from the electrode for 3 minutes.

We were then able to use multiple electrophysiological measurements and analyses of single and multi-unit spiking activity as well as local field potentials to determine the location of the silicon probe relative to the laminar structure of the hippocampus. Figures 3 and 4 illustrate these techniques. In these figures, the recorded 31 intra-hippocampal channels (200 μm spacing) are, in turn, aligned to a coronal section of the dorsal hippocampus stained with LFB/CV from the same animal. Alignment of the probe relative to the histology was based on recovered artifact of the probe tips (not shown) and stereotactic coordinates used during insertion. Vertical alignment of histology and electrophysiology depicted in Figures 2 and 3 are only approximate as the plane of insertion was oblique to the histological coronal plane and the relative thicknesses of the hippocampal layers change along the AP axis. In one illustrative animal, depicted in Figure 3, isolation of single unit spiking activity on all channels revealed a large cluster in the region of stratum pyramidale and a single channel in the region of stratum granulosum. In four animals, the position of the pyramidal cell layer was confirmed by calculating multi-unit spiking activity power in the 600-6,000 Hz frequency band (Figure 3D, mean ± SEM, n = 4). The traces from these animals were aligned at the peak power of the 600-6,000 Hz band, representing the pyramidal cell layer.

Several analyses of the local field potentials also confirm precise localization of the probe within the hippocampus (Figure 4). We first examined the wide-band local field potentials across all 31 intra-hippocampal channels (Figure 4A). Inspection demonstrates prominent multi-unit spiking activity in a band near the dorsal end of the electrode and a less-prominent band of spiking activity more ventrally (Figure 4A), which is also reflected in the isolated single unit activity (Figure 3B and 3D). We also consistently observed waves of synchronized activity in stratum radiatum with a frequency of 2-3 Hz (Figure 4A). These waves demonstrated a phase reversal at the pyramidal layer, at the bottom of stratum radiatum reminiscent of sharp waves described in the awake rodent hippocampus (Bragin et al., 1995b; Buzsáki et al., 2003), and again at dentate gyrus middle molecular layer (arrows). It is possible that these are regular sharp-wave-like (SPW-like) events representing synchronized pre-synaptic currents from CA3 projections, as SPW have been shown to take on an oscillatory characteristic under isoflurane anesthesia in rodents (Lustig et al., 2015).

We also looked at the spectral power across the laminar structure of the porcine hippocampus in several frequency bands that are prominent in the rodent hippocampus (Bragin et al., 1995a), including classic theta (θ, orange trace, 4 - 10 Hz), low gamma (γlow, blue trace, 25 - 55 Hz), high gamma (γhigh, purple trace, 55 - 90 Hz), and ripple range (ρ, red trace, 150 - 250 Hz) bands (Figure 4B, mean ± SEM, n = 4). Oscillations within a classic theta frequency band (orange trace) appear prominently in stratum radiatum with a sharp fall-off towards the hippocampal fissure, whereas theta oscillations in the rodent demonstrate increasing power along the depth of CA1, peaking in stratum lacunosum-moleculare (Buzsáki et al., 2003). This discrepancy could represent a difference in species, or an effect of anesthesia. Low and high gamma bands (Figure 4B, blue and purple traces) also have peaks of power in stratum radiatum as well as in stratum pyramidale. Since the traces from these four animals were aligned at the peak power of the 600-6,000 Hz band (the pyramidal cell layer), it left 25 channels for the intra-hippocampal laminar recordings, and excluded the channels at the bottom of the probe positioned in the dentate gyrus layer (Figure 4B, also see Figure 3D).

Discrete oscillatory events within the frequency range of 150 Hz – 250 Hz have been described in the pyramidal cell layer of the awake and asleep rodent hippocampus, termed “ripples” (Buzsáki, 2015). Although it is thought that ripple events in the LFP arise from a coordinated interaction between pyramidal cells and local interneurons, high frequency oscillatory events in the pyramidal cell layer in general may have a significant component of their measured power derived from synchronized pyramidal cell action potentials. Importantly, much of the power may come from action potentials of cells further than 100 μm away, making this signal detectable even when there are no neurons close enough to a recording electrode to result in isolatable single unit activity (Schomburg et al., 2012). This phenomenon may be particularly relevant in the current study as we did not observe any definitive ripple oscillatory events in stratum pyramidale, most likely due to effects of anesthesia. Figure 4B shows integrated spectral power in this ripple band (150 Hz – 250 Hz) across channels (red trace), demonstrating a large peak at the level of the prominent multi-unit spiking activity, which is also the channel of maximum power within the 600 - 6,000 Hz band (Figure 3D), suggesting localization to the CA1 pyramidal layer. To further describe porcine hippocampal layers, we performed perforant path stimulation in one of the animals. The responses to the perforant path stimulation were averaged over 50 msec time period (20 stimulations, 2 seconds apart). The laminar profile of the evoked potentials is shown in Figure 4C. Two phase reversals were observed, above and below stratum radiatum hippocampal layer (Figure 4C, arrows). Localized damage from the insertion of the electrode through the pyramidal CA1 cell layer was confirmed post mortem with LFB/CV staining (arrow, Figure 3C). Additionally, these electrode tracks were confirmed with APP labeling, which accumulates in injured axons (Gentleman et al., 1993; Johnson et al., 2013; Sherriff FE, 1994) and may be useful for electrode track detection in acute experiments in other species (data not shown).

## Discussion

We describe here aspects of the laminar structure of the dorsal hippocampus of the Yucatan miniature pig both histologically and electrophysiologically using a methodology of stereotaxis that does not require individual subject imaging. This is important as pigs have emerged as an attractive large animal model for neuroscience research and translational CNS disease models (Gieling et al., 2011b; Lind et al., 2007), yet little has been reported about the comparative laminar structure of the hippocampus and its electrophysiology. Due to their gyrencephalic brains and substantial white matter, pigs have been used to model CNS disorders such as TBI (Cullen et al., 2016) and epilepsy (Van Gompel et al., 2011), and may be useful for other hippocampal centric models of cognitive dysfunction (Maas et al., 2017; Rollin, 2006). Further, these models are strengthen by the growing number of studies conducted to measure cognitive performance in pigs (Asher et al., 2016; Broom et al., 2009; Conrad and Johnson, 2015; Fleming and Dilger, 2017; Gieling et al., 2011a; Gieling et al., 2011b; Grimberg-Henrici et al., 2016; Schramke et al., 2016; Schuldenzucker et al., 2017; Sullivan et al., 2013).

Awake electrophysiological recordings in pigs have been reported in the past, and the use of pigs for intracranial monitoring studies and *ex vivo* electrophysiology has increased recently (Conn, 1991; Van Gompel et al., 2011; Wolf, 2017). One barrier to acute or chronic electrophysiology of the hippocampus has been the lack of methodology for targeting the hippocampus precisely without imaging guided stereotaxis. While several stereotaxic instruments have been developed for pigs, variations of skull landmarks due to age-depended changes can create inconsistency in stereotaxic procedures (Ettrup et al., 2011; Marcilloux et al., 1989; Poceta et al., 1981; Saito et al., 1998; Szteyn et al., 1980). We have therefore developed an inexpensive, non-imaging based swine stereotaxis for *in vivo* electrophysiology, which is based on skull position, angle and landmarks and then refined with electrophysiological mapping. Localization within the dorsal hippocampus needs to be determined more precisely as well as independently for each subject when localization of layers is required for recordings or implantation (Saito et al., 1998). In order to accomplish this, we implemented electrophysiological mapping based on DBS techniques widely used for neurosurgical procedures (Gross et al., 2006). This allows us to target multichannel electrodes to the pyramidal CA1 cell layer and other hippocampal laminar structures during insertion, and to reduce the reliance upon histological determination of suitable electrode placement within the hippocampus. While intra-operative recording to verify placement of chronic electrodes is not a standard procedure even in chronic primate electrophysiology, we suggest that it may complement existing image guided techniques as more complex laminar probes begin to be utilized in large animals.

Our histological investigation of the dorsal hippocampus of the Yucatan miniature pig confirms a preserved laminar structure across species with a standard orientation of the hippocampal layers. Our cytoarchitectural findings are consistent with a previously published, extensive immunohistological characterization of cell types in porcine hippocampus, including the progressive dispersion of pyramidal cells as one approaches the subiculum (Holm et al., 1992; Holm et al., 1993; Holm et al., 1990; Holm and West, 1994). Interestingly, we found that this variability in pyramidal cell density was reflected in the electrophysiology. Where the pyramidal cells were spread out, the peak power of the measured spiking activity was broad and less prominent than might be observed in the rodent (see Figure 2H). Further, as seen in Figure 1, this dispersion juxtaposes the pyramidal cell bodies of some cells with the dendritic arbors of others, potentially blurring the distinct layering of input and output currents (and thus the patterns of sinks and sources) classically seen in awake rodent electrophysiological studies. We also report the thickness of each hippocampal layer and directly compare them to those of the rat hippocampus. Porcine hippocampal layers including the alveus and strata pyramidale, radiatum, lacunosum-moleculare, and moleculare were significantly larger than those in rats, while other layers, namely strata oriens and granulosum, were not significantly different. This difference could reflect a disproportional increase in inputs from afferent cortex relative to local intra-hippocampal circuitry as the gyrencephalic neocortex increases in size and complexity along the phylogenetic tree, while basic hippocampal function presumably remains largely preserved.

We have simultaneously recorded electrophysiological activity of many individual cells in the porcine dorsal hippocampus, which are appropriately distributed in a laminar fashion as expected from the histology and the LFP structure. The stereotaxic locations of the cell body layers were confirmed with a power analysis of the 600 – 6,000 Hz frequency bands, which identifies multi-unit spiking activity corresponding to strata pyramidale and granulosum. Importantly, our spectral power analysis of the ripple band (150Hz – 250 Hz) also identified the pyramidal cell layer (see Figure 4B). As discussed above, power in this band may have significant contributions from synchronized pyramidal cell action potentials, including from cells over 100 μm from the recording electrode (Schomburg et al., 2012), a distance too great to routinely pick up spiking potentials. This could allow electrophysiological identification of the pyramidal layer in circumstances without detectable spiking activity, such as under deep anesthesia, or when only macro/low impedance electrodes are utilized.

In addition to the ripple frequency band, we also looked at the power of several other physiologically relevant bands expected in the hippocampus based on rodent experiments. Classic theta in the awake rodent hippocampus (4 Hz – 10 Hz) shows a spectral power laminar pattern that increases from stratum radiatum to the hippocampal fissure, where it peaks, decreasing in deeper lamina (Bragin et al., 1995a). In contrast, we observed a broad peak in theta power in the stratum radiatum. Gamma oscillation power (25 Hz – 100 Hz) in the awake rodent hippocampus shows a persistent increase from the stratum radiatum down to the hilus (Bragin et al., 1995a). Again, in contrast, we see only a broad peak in gamma power (25 Hz – 55 Hz and 55 Hz – 90 Hz) in stratum radiatum, with an additional peak in the pyramidal cell layer in the high gamma band. There could be a number of reasons for these discrepancies. First, the relative strengths and patterns of input into the pyramidal cell dendritic arbor may be fundamentally different in pigs compared to rats. This could result either in completely different laminar patterns, or in a shift in the physiologically relevant frequency bands. An alternative or complementary explanation may be that the pattern and strength of inputs arriving into the dendritic arbor of the CA1 pyramidal cells are likely affected by the depth of anesthesia (Lustig et al., 2015).

There is also evidence that adequate depth of isoflurane anesthesia can drive the rodent hippocampus into a SPW-like state with bursts of inputs into stratum radiatum that demonstrate a similar pattern to CA3-derived SPW observed in the awake rodent (Lustig et al., 2015). We also observe regular bursts of synchronized input into stratum radiatum in the isoflurane anesthetized pig that have the typical phase reversal across the pyramidal cell layer expected of CA3-derived SPWs. Consistent with the anesthetized state in the rodent (Lustig et al., 2015), we did not observe discrete ripple events, nor as prominent a ripple oscillation as those reported in behaving animals, but a peak in the ripple band frequency (150 – 250 Hz) in stratum pyramidale was present under anesthesia and correlates with the laminar profile of spiking activity (Buzsáki, 1989).

Our efforts to characterize network disruptions after TBI illustrate how the pig can also be used as a translational model of neurological disorders, taking advantage of this characterization of the pig hippocampus and its electrophysiology. Electrophysiological changes have largely been studied in rodent models of TBI, including disruptions in long-term potentiation, synaptic plasticity and broadband power (Miyazaki et al., 1992; Paterno et al., 2015; Schmitt and Dichter, 2015; Villasana et al., 2015; Zhang et al., 2011). In a pig TBI model we have observed hippocampal axonal and synaptic dysfunction as well as regional hyperexcitability in an *ex vivo* slice preparation following an established porcine model of TBI where diffuse axonal injury is the primary neuropathological finding (Cullen et al., 2016; Johnson et al., 2016; Meaney et al., 1995; Wolf, 2017). Importantly, the electrophysiological characterization and methodology described here will permit the confirmation of these results in anesthetized and awake behaving swine using an *in vivo* electrophysiological procedure which utilizes laminar multi-electrode silicon probes. This will further support investigations into hippocampal-centric translational models such as those being developed for the study of post-traumatic epilepsy, as well as laying the groundwork for closed loop neuromodulation-based therapies in these models.

To summarize, we have characterized porcine laminar hippocampal electrophysiology under anesthesia utilizing a new laminar probe and using techniques that don’t require imaging-based stereotaxis. We have histologically confirmed that the neuroanatomy of Yucatan porcine brain closely resembles previously reported hippocampal structure, with features of both rodent and primate hippocampus. We have precisely placed depth probes into the dorsal hippocampus of miniature pigs, and developed electrophysiology-based methodology to identify hippocampal layers and to record hippocampal single units and local field potentials simultaneously across these layers using multichannel probes. This will allow for analysis of spike-field interactions during normal and disease states. Laminar structure of hippocampus and the precise location of custom multichannel silicon probes were confirmed by electrophysiology and histopathology, reducing the future reliance upon immediate recovery of electrode tracks. We hope that this characterization of the miniature swine hippocampus and stereotactic localization of the dorsal hippocampus will allow for greater use of these gyrencephalic mammals in translational models of hippocampal circuitry, as well as awake neurophysiology under healthy and pathophysiological conditions such as TBI and associated epilepsies. The combination of complex behaviors generated by these animals and the detailed neurophysiology of the hippocampus may present an ideal “middle ground” for translational hippocampal-centric disease models prior to clinical trials.

